# A viroid-like RNA can be transmitted among different *Trichoderma* species affecting their antagonistic capacity

**DOI:** 10.64898/2026.01.28.702247

**Authors:** Cristina Formiglia, Marco Forgia, Beatriz Navarro, Francesco Di Serio, Nadia Serale, Safa Oufensou, Virgilio Balmas, Quirico Migheli, Niccolò Miotti, Olga Rueda, Federica Bono, Marcos de la Peña, Massimo Turina

## Abstract

Viroids are small, circular non-coding RNAs that autonomously replicate in plants, exploiting host cellular machinery for replication and spread. Recent studies reveal that viroid-like agents can infect filamentous fungi, suggesting cross-kingdom interactions. In this study, we report the discovery and the characterization of TsvlRNA1 in *Trichoderma spirale*, a transmissible viroid-like RNA containing a hammerhead ribozyme in one polarity strand. Bioinformatic data, molecular validation, and reverse genetics experiments demonstrate that TsvlRNA1 is circular with an active ribozyme essential for replication. TsvlRNA1 replicates autonomously and transmits horizontally between *Trichoderma* species, eliciting 21–23 nt viroid-derived small RNAs consistent with RNA silencing targeting. The biocontrol capacity of *Trichoderma* against *Rhizoctonia solani* is variably modulated by TsvlRNA1, with effects ranging from positive to negative depending on host strain. In *T. spirale*, data suggests genotype-by-agent interactions influence antagonistic potential negatively. TsvlRNA1 transmission via horizontal routes is prevalent, and the viroid-like RNA fails to infect plant hosts experimentally. These results highlight so-far the underappreciated ecological and functional diversity of viroid-like agents in fungi, with implications for fungal biology, biocontrol, and genotype-phenotype relationships in eukaryotes.

**Importance:** Species of the fungal genus *Trichoderma* play a central role in sustainable agriculture by controlling fungal plant pathogens and supporting plant growth. For this reason, *Trichoderma*-based products represent a substantial share of the global market for microbial biofungicides. Viroids are the smallest known infectious agents, and their presence in filamentous fungi has only recently been discovered. Consequently, little is known about their biology, transmission, or interactions with fungal hosts. In this study, we describe TsvlRNA1, a viroid-like RNA associated with *T. spirale*, representing only the second viroid-like RNA to be biologically characterized in fungi. We show that TsvlRNA1 can influence the ability of *Trichoderma* to inhibit *Rhizoctonia solani*, a major plant pathogen, demonstrating its biological relevance. Unexpectedly, TsvlRNA1 can be transmitted between different *Trichoderma* species. This finding raises concerns about the possible transfer of genetic traits between fungi, including those related to fungicide resistance, with important implications for agricultural biocontrol.

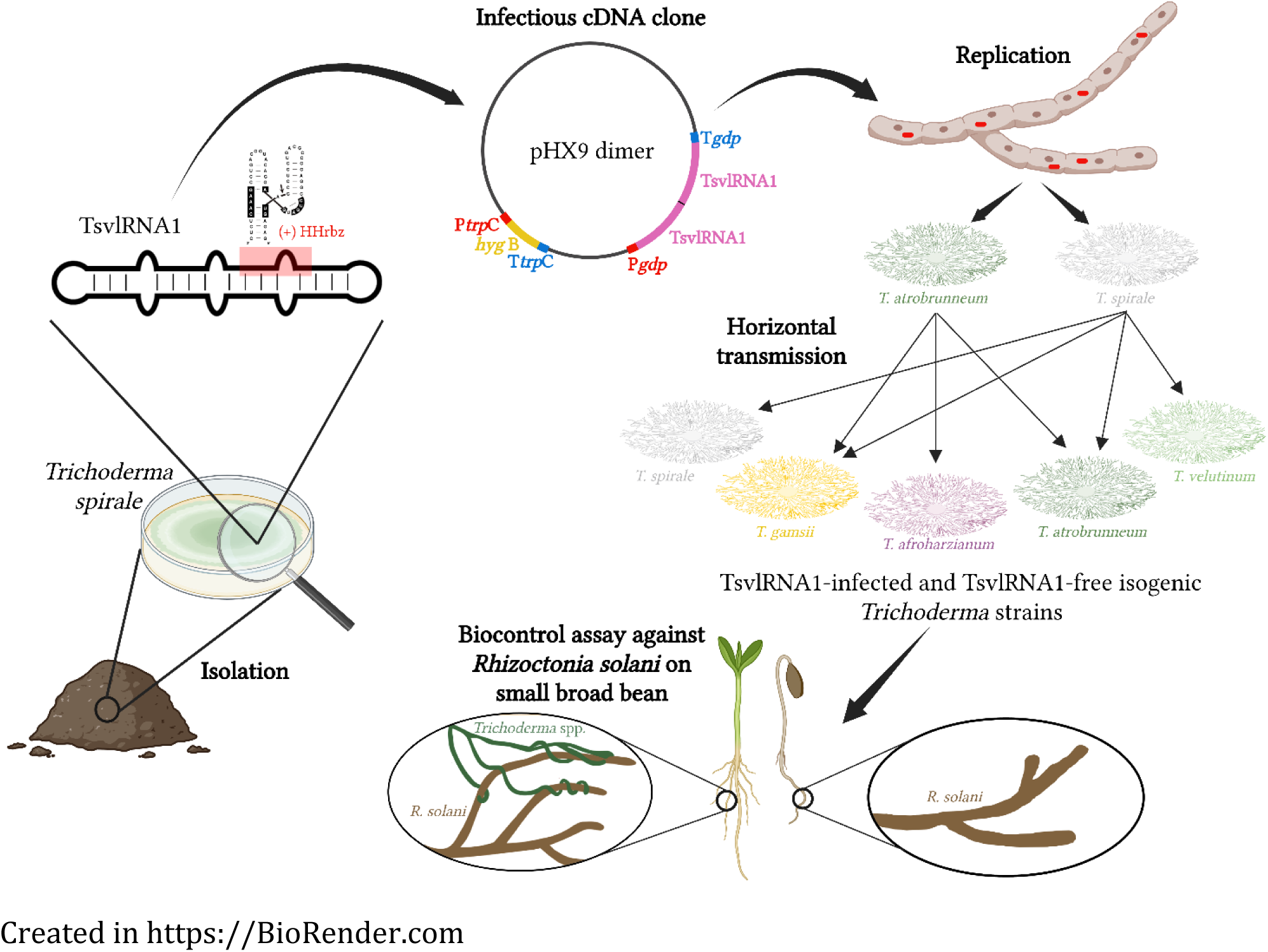

## Introduction

Viroids, first identified as plant pathogens in the early 1970s (1), are recognized as the smallest replicators. They consist of a covalently closed circular (ccc) RNA ranging from 234 to 506 nucleotides in length, exhibiting high self-complementarity, thus assuming a rod- or branch-shaped conformation (2–4). Viroids lack protein-coding capacity and some of them contain a self-cleaving ribozyme in each polarity strand that catalyzes a site-specific cleavage of viroid oligomeric RNAs during replication (5–8).

Viroids are classified into two families (6,8): *Pospiviroidae*, which groups viroids replicating and accumulating within the nucleus, and *Avsunviroidae*, which includes viroids that replicate and accumulate in chloroplasts and possess hammerhead ribozymes (HHrbz) (9,10).

Replication occurs through a rolling-circle replication (RCR) mechanism comprising three steps: RNA transcription, processing, and ligation. *Pospiviroidae* follow an asymmetric RCR pathway, catalyzed by host enzymes such as RNA polymerase II, RNase III-like, and DNA ligase I (2,8). In contrast, *Avsunviroidae* follow a symmetric RCR pathway mediated by the nuclear-encoded and plastid localized RNA polymerase, self-cleaving ribozymes, and the chloroplast tRNA ligase (11,12). While in the symmetric RCR pathway, both genomic and antigenomic RNAs are cleaved and circularized, in the asymmetric pathway only genomic RNAs undergo these steps (2,13,14).

Viroid-like RNA (vdlRNA) elements include plant virus satellite RNAs, retroviroids, retrozymes and other vdlRNA elements with larger genomes and protein-coding capacity (Ribozyviruses) (15–19). A example is the hepatitis delta virus, a human pathogen that encodes the delta antigen (20). Recently, a diverse array of viroid-like agents, including ambiviruses, some mitoviruses, Zeta viruses, and Obelisks, has been characterized, extending the host range of vdlRNAs to fungi and bacteria (5,21–23).

The fungal genus *Trichoderma* (teleomorph *Hypocrea*, Hypocreales, Ascomycota) is widely distributed across ecosystems, including agricultural fields and decaying plant material (24). Nearly 500 species have been identified globally (25), many of which are exploited for biocontrol of fungal phytopathogens through mechanisms like competition, antibiosis, mycoparasitism, induction of plant defense responses, and promotion of plant growth (26). Additionally, *Trichoderma* species are utilized in the production of antibiotics, enzymes, and biofuel (27–29) and in the bioremediation of xenobiotic compounds in water and soil (30).

Fungal cytoplasmatic infectious elements (mycoviruses and vdlRNAs) are transmitted horizontally within populations via hyphal fusion. A barrier to hyphal fusion is the vegetative incompatibility (VIC), which regulates self/non-self recognition through *het/vic* genes and limits horizontal mycovirus transmission (31). Compatibility among *Trichoderma* isolates has been examined via heterokaryon —structures in which genetically distinct nuclei coexist within a common cytoplasm— formation through hyphal anastomosis or protoplast fusion (32,33). However, most studies precede molecular techniques, which expanded the genus to 459 species (34). Recent analyses of vegetatively compatible groups (VCG) in *Trichoderma* are scarce and largely based on morphological observations (35).

This study investigates the presence of a vdlRNA sequence in a previously characterized collection of *Trichoderma* spp. isolates from Sardinian uncultivated soils (36,37). We characterized a vdlRNA isolated from *T. spirale*, previously detected as ORFan segment of putative viral nature (36). An infectious clone was successfully developed, enabling demonstration of its autonomous replication and horizontal transmission to different *Trichoderma* species following co-culture transfection. This achievement allows for the first time to prove the role of a vdlRNA in influencing the antagonistic properties of *Trichoderma* spp.

## Methods

### Origin of fungal isolates, ORFan/ribozyme analysis of the Sequence Read Archive, and in vitro culture on potato dextrose agar

The fungal isolates used in this study, belonging to the genus *Trichoderma*, are part of a collection housed at the University of Sassari, Italy, and assembled since 2009 (37). An ORFan identified in the study by Pagnoni et al. (36) was further investigated using the INFERNAL algorithm on Serratus cloud computing platform (5), designed for the detection of circular molecules and ribozymes. The strain *T. spirale* (T45) found to carry the viroid-like element was subjected to Next-Generation Sequencing analysis (Bioproject PRJNA1303385, Accession Number SRR34918256).

A viroid-free isogenic *T. spirale* (T45neg) colony was obtained by preparing a conidial suspension from TsvlRNA1-positive T45 and selecting physically isolated colonies, which were screened for the absence of vdlRNA.

Isolates used in this study were placed on Potato Dextrose Agar (PDA; Sigma-Aldrich, St. Louis, MO, USA) medium and incubated at 28°C for 8-10 days under near-ultraviolet light to induce conidiation.

### Identification of different *Trichoderma* species

Sequence data from the internal transcribed spacer (ITS) regions, the RNA polymerase II subunit (RPB2), and the translation elongation factor 1-α gene (TEF1-α) were used to perform a multi-locus molecular phylogenetic analysis, thus identifying the *Trichoderma* species.

Genomic DNA from each strain was extracted from fresh mycelium, following the thermolysis method described by Zhang et al. (38).

PCR amplification of the ITS, RPB2, and TEF1-α gene fragments was performed using OneTaq DNA polymerase (NEB, USA) and gene-specific primers (Table S1). Amplicons were purified with the ‘Zymo gel DNA recovery kit’ (Zymo research, Irvine, CA, USA), and bidirectionally sequenced by Biofab Research (Rome, Italy).

For species identification, ITS, RPB2, and TEF1-α sequences from isolates T22, T36, T45, T71, TO71B, T84, T99, and T100 were aligned with reference sequences (Table S2). MUSCLE alignment was performed in MEGA11, and a maximum-likelihood phylogenetic tree was generated with IQ-TREE (39), with clade support assessed by 1000 bootstrap replicates using *Protocrea pallida* as the outgroup.

### Transformation vector assembly

Inverse PCR with primers CHV1-25Rev and CHV1-12700-For (Table S1) released the CHV1 genome from the transformation plasmid pHX9 (40), generating pHX9-nv with multiple cloning sites under the *Cryphonectria parasitica gpd-1* promoter/terminator, and carrying the *Escherichia coli hygB* selectable marker.

The expression vector ‘pHX9 GFP’ was constructed to transgenically express green fluorescent protein (GFP) by inserting the GFP sequence —amplified from clone R3-p21-TripB-GFP (41) using primers introducing *NotI* and *StuI* sites— into *NotI*-*StuI*–digested pHX9-nv.

RNA from *T. spirale* (T45) naturally infected with TsvlRNA1 was reverse transcribed with random primers. Two full-length genomic fragments of TsvlRNA1 (A and B) were PCR-amplified (primers in Table S1), cloned into pCR™-Blunt vector (Invitrogen™, Carlsbad, CA), excised with *EcoRI*/*NotI* (A) and *EcoRI*/*KpnI* (B), and ligated into *KpnI*/*NotI*–digested pBluescript vector using T4 DNA Ligase (Thermo Fisher Scientific, Waltham, MA, USA) to generate plasmid ‘pBlu-Trichodimer#21’. The dimer sequence was verified by sequencing, then inserted via *NotI*/*KpnI* into pHX9-nv to create ‘pHX9 dimer’ (Fig 1).

**Figure 1.**
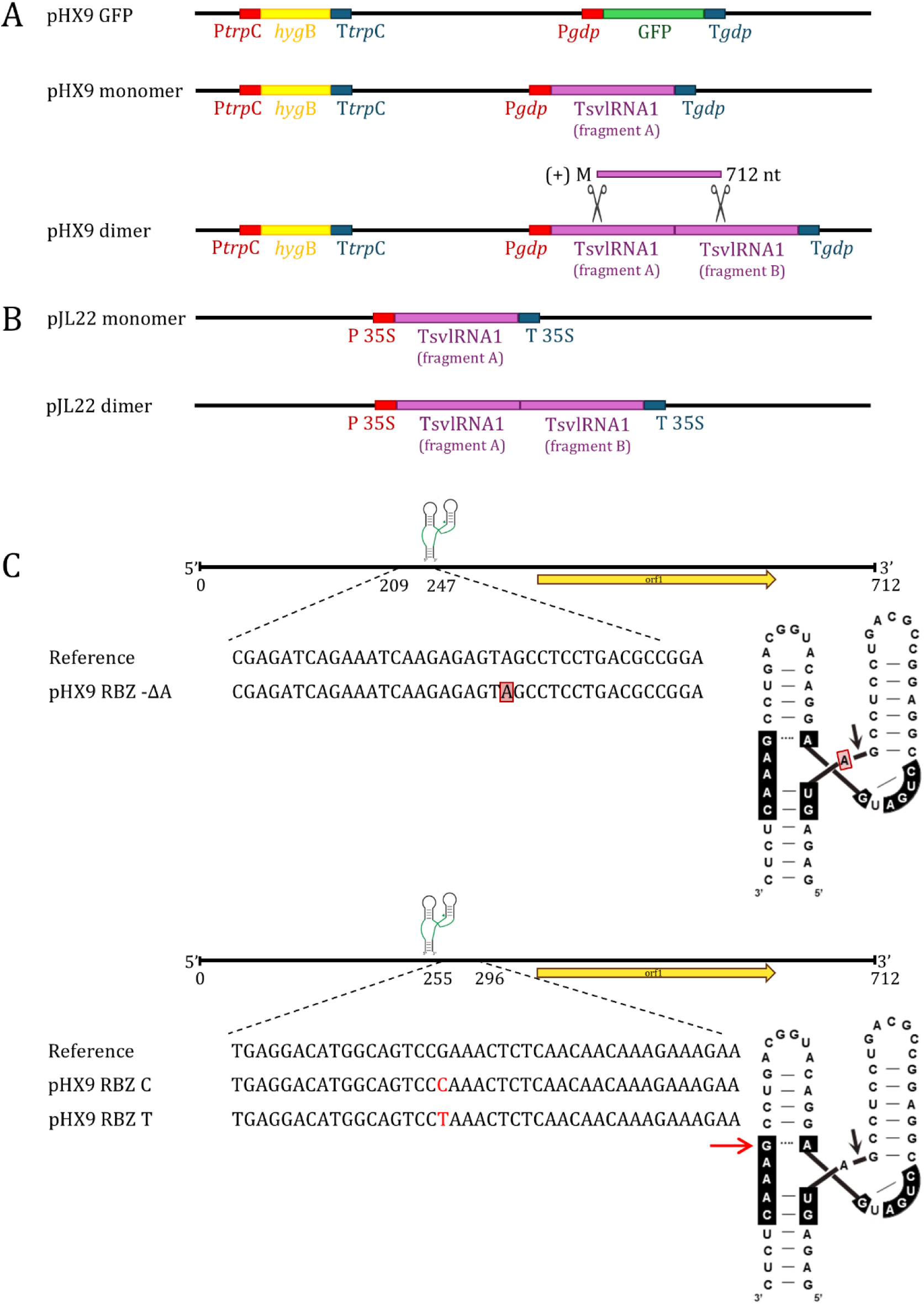
Schematic representation of recombinant plasmids used in this study. The cloning operations used to construct individual expression vectors are described in detail in the materials and methods section. **A** Plasmid backbone named pHX9-nv contains *Escherichia coli* hygromycin B phosphotransferase gene (*hygB*) as a selectable marker, flanked by the transcriptional control elements of the *trpC* gene from *Aspergillus nidulans*. GFP gene under the control of *gpd-1* gene promoter and terminator from *C. parasitica* were introduced creating ‘pHX9 GFP’. Infectious cDNA clones of TsvlRNA1 were generated introducing the monomeric sequence and the dimeric sequence of TsvlRNA1, respectively, flanked by the same promoter and terminator from *C. parasitica*. They are called ‘pHX9 monomer’ and ‘pHX9 dimer’. The two scissors indicate the position of the hammerhead ribozyme present on the positive strand; its self-cleavage activity generates the (+) monomer, which is 712 nt in length. **B** pJL22 plasmid was used to generate two constructs: one containing the monomeric sequence of TsvlRNA1 and the other the dimeric form, named as ‘pJL22 monomer’ and ‘pJL22 dimer’. Both sequences are flanked by the Cauliflower Mosaic Virus 35S promoter and terminator. **C** Recombinant clones derived from ‘pHX9 dimer’ with various mutations targeting the (+) HHrbz active site include a deletion of adenine at position 231 (‘pHX9 RBZ-ΔA’) and point mutations substituting cytosine or thymine at position 272 (‘pHX9 RBZ C’ and ‘pHX9 RBZ T’, respectively). The deletion is indicated by the red box. In panel C only one of the two monomers is displayed to allow a more in-detail view of the mutations performed in both monomers constituting the dimer clone.

To create ‘pHX9 monomer’, the pCR™-Blunt vector (Invitrogen™, Carlsbad, CA) containing fragment A was digested with *Not*I*/Kpn*I and the excised sequence was inserted into pHX9-nv (Fig 1).

Three TsvlRNA1 mutants —deletion of adenine (A) at position 231 and cytosine (C) or thymine (T) substitutions at position 272— were generated by inverse PCR (Table S1) and cloned as described to yield plasmids ‘pHX9 RBZ -ΔA’, ‘pHX9 RBZ C’, and ‘pHX9 RBZ T’ (Fig 1). All the mutated fragment A and B were Sanger sequenced before assembling the dimer.

### *Trichoderma* sensitivity test to hygromycin B and protoplasts transformation

The minimum inhibitory concentration (MIC) of hygromycin B required to inhibit *T. lixii*, *T. spirale*, and *T. atrobrunneum* (T36, T45neg, T99) growth was determined on PDA with increasing concentrations and used to select transformants.

Protoplasts derived from T36, T45neg, and T99 were produced and transformed according to Malmierca et al. (42). Transformants were sequentially transferred to medium with twice the MIC of hygromycin B, then to non-selective and finally to selective medium to obtain mitotically stable isolates, which were grown on cellophane-covered PDA for 4–5 days before mycelia were harvested for RNA extraction.

### GFP visualization

Mycelium from ‘pHX9 GFP’ transformants was transferred onto glass slides. GFP fluorescence was captured with a Zeiss LSM 900 confocal microscope (488-nm excitation, 500–525-nm emission) and analyzed with ZEN 2.3 software (Carl Zeiss AG, Oberkochen, Germany).

### RNA extraction, DNase treatment, cDNA synthesis and quantitative RT-qPCR

RNA extraction was carried out using the ‘Spectrum Plant Total RNA Kit’ (Sigma-Aldrich, St. Louis, MO, USA), in accordance with the manufacturer’s instructions. RNA concentration was measured using a Nanodrop LITE Spectrophotometer (Thermo Fisher Scientific, Waltham, MA, USA). DNase treatment using ‘DNA-free™ DNA Removal Kit’ (Ambion™, Austin, TX, USA) preceded cDNA synthesis using the ‘High-Capacity cDNA Reverse Transcription Kit’ (Thermo Fisher Scientific, Waltham, MA, USA) with random primers; finally, cDNA was diluted 1:5.

Minus-strand cDNA was generated using a TAG-containing primer (TrichoRBZ TAG NEGS in Table S1) and then purified with the ‘Zymo DNA Clean & Concentrator®-25 kit’ (Zymo Research, Irvine, CA, USA).

Primer design was conducted using NCBI primer BLAST tool (https://www.ncbi.nlm.nih.gov/tools/primer-blast/); resulting primers are listed in Table S1. Minus strand-specific and total cDNA-specific Real-Time qPCR (RT-qPCR) with TsvlRNA1-specific primers/probe were performed on DNase-treated/untreated RNA and corresponding cDNA to verify transgene integration and assess vdlRNA replication. RT-qPCR assays were performed in 10 μL using iTaq™ Universal Probes Supermix (BioRad, Hercules, USA) on a CFX Connect RT-qPCR Detection System (Biorad, Hercules, USA).

### Analyses of RNA self-cleavage in vitro and 5’ RACE

The *NotI*- or *KpnI*-linearized ‘pBlu-Trichodimer#21’ plasmid served as the template for T7 or T3 in vitro transcription of TsvlRNA1 dimeric transcripts of plus and minus polarity, respectively. Transcripts were analyzed by denaturing 5% PAGE containing 8 M urea and 1X TBE, and products were recovered for self-cleavage assays and 5′ RACE. TsvlRNA1 primary transcripts were resuspended in Tris-HCl buffer at pH 7.5 or 8.5, denatured at 95 °C, gradually cooled to 37 °C or 25 °C, and then incubated with different MgCl₂ concentrations to induce self-cleavage. For 5′ RACE, the eluted RNA was reverse-transcribed with ‘Superscript IV’ (Invitrogen™, Carlsbad, CA) and primer RACEpos349 (Table S1), poly(dG)-tailed, PCR-amplified, cloned and sequenced according to Hirtzmann methodology (43).

### Northern blot hybridization assays

DNase-treated, denatured RNA was resolved on denaturing 5% PAGE containing 8 M urea and 1X TBE, electroblotted onto nylon membranes, UV-crosslinked, and hybridized with DIG-labelled riboprobes specific for (+) or (−) TsvlRNA1 strands. Riboprobes were synthesized using T7 transcription with the DIG-RNA Labelling Mix (Roche Diagnostics GmbH, Germany) from *SpeI*-linearized pGEM-T Easy containing a partial TsvlRNA1 insert in the correct orientation. Pre-hybridization and hybridization were performed in DIG Easy Hyb (Roche Applied Science, Germany) at 65 °C, and signals were detected using anti-DIG alkaline phosphatase antibody fragments and the chemiluminescent substrate CDP-Star (Roche Applied Science, Germany) on a ChemiDoc Touch system (Bio-Rad, Hercules, CA, USA).

### Horizontal and vertical transmission and extracellular acquisition of TsvlRNA1

Donor strains of *T. spirale* (T45DIM5) and *T. atrobrunneum* (T99DIM3) expressing the ‘pXH9 dimer’ were co-cultured with different recipient strains to evaluate whether transgene expression was required for TsvlRNA1 replication (all combinations are listed in Table 1). Donor and recipient strains were placed 40 mm apart on PDA, and recipient plugs collected after 6 and 11 days were subcultured and screened for hygromycin sensitivity. TsvlRNA1 horizontal transmission was then assessed by RT-qPCR and northern blot assays. The same combinations were tested using TsvlRNA1-negative donors (T45neg, T99) to examine interaction dynamics. Vertical transmission was assessed from conidia of TsvlRNA1-positive *T. atrobrunneum* isolates (T84ANAS-1, T99ANAS-1) obtained through co-culture.

**Table 1.** Table showing all combinations tested in horizontal transmission experiments between dimer-transformed viroid-like *Trichoderma spirale* (T45DIM5) and *Trichoderma atrobrunneum* (T99DIM3) isolates and viroid-like-free recipient isolates. ‘X’ indicates absence of TsvlRNA1 transmission, while ‘✓’ indicates successful transmission.

| Donor | Recipient |  |  |  |  |  |  |  |
| --- | --- | --- | --- | --- | --- | --- | --- | --- |
|  | T22 | T36 | T45neg | T71 | T071B | T84 | T99 | T100 |
| **T45DIM5** | X | X | ✓ | ✓ | X | ✓ | X | ✓ |
| **T99DIM3** | ✓ | X | X | ✓ | ✓ | ✓ | ✓ | X |

To test for possible extracellular acquisition of the viroid-like RNA, T99 mycelia were collected from liquid culture, homogenized in distilled water, washed with STC buffer (1 M sorbitol, 10 mM Tris·HCl, pH 7.5, 20 mM CaCl2), and recovered by low-speed centrifugation. Hyphal fragments were mixed with total RNA from T45ANAS-A or in vitro produced RNA, incubated on ice and at room temperature for 30 minutes each, and then grown at 28 °C for 4–5 days in liquid medium and recovered for RNA extraction.

### Small RNA sequencing

The small RNAs (sRNAs) of four TsvlRNA1-positive *Trichoderma* isolates —including two also infected by mycoviruses—were sequenced to assess silencing-related antiviral responses (T22ANAS-1, T45ANAS-A, T71ANAS-A, and T99ANAS-1). sRNA libraries were prepared and sequenced on an Illumina NovaSeq X by Novogene Bioinformatic Technology (Beijing, China). The raw sequencing reads were deposited in the NCBI Sequence Read Archive under the BioProject PRJNA1301113 (Accession Number SRR34855465, SRR34855464, SRR34855463, and SRR34855462). After trimming with Trimmomatic (15–50 nt) (44), >10 million reads per sample remained. Reads were mapped to TsvlRNA1 and viral genomes with BWA (45), allowing circular genome alignment. Sorted BAM files were filtered with SAMtools (46), and strand-specific and size-distribution analyses were generated and visualized with Tablet (47).

### Evaluation of *Trichoderma* antagonistic activity against fungal infection in small broad bean

Agar plugs bearing mycelium of the soilborne polyphagous pathogen *Rhizoctonia solani* J.G. Kühn (AG-4), previously isolated from quinoa (*Chenopodium quinoa* Willd.), were excised from 7-day PDA cultures and placed onto each *Trichoderma* mycelial plug; both were placed in a seedbed containing sterilized potting mix (Vigorplant Italia). After one-week, small broad bean (*Vicia faba* var. *minor* Beck) seeds were sown, and the seedbeds were incubated in a glasshouse for 21 days under controlled temperatures and daily irrigation. For each *Trichoderma* isolate, three replicates of nine plants were prepared. After three weeks, the disease severity index was assessed using a five-class empirical scale (0–4). A completely randomized design was used, and data were analyzed via one-way ANOVA with Honestly Significant Difference and by Kruskal–Wallis with Dunn’s post-hoc test.

### TsvlRNA1 viroid-like expression in plants

Transient expression of TsvlRNA1 in plants was achieved by cloning its monomeric and dimeric genomic sequences into the binary vector pJL22 (48) under the Cauliflower mosaic virus 35S promoter and terminator. Monomer and dimer inserts were excised from pBlu-Trichodimer#21 using *EcoRI*/*KpnI* or *KpnI*/*NotI*, respectively, purified with the ‘Zymoclean Gel DNA Recovery Kit’ (Zymo Research, Irvine, CA, USA), and ligated into linearized pJL22 to generate the constructs ‘pJL22 monomer’ and ‘pJL22 dimer’. Each construct was used to transform *Agrobacterium tumefaciens* strain C58C1, which was grown on selective medium and resuspended in MES buffer for infiltration.

One-month-old *Nicotiana benthamiana* plants were agroinfiltrated with *A. tumefaciens* carrying ‘pJL22 dimer’ or ‘pJL22 monomer’, together with clones expressing the silencing suppressor p19 and, in some treatments, the tobacco mosaic virus (TMV) movement protein. Plants were maintained for one week under controlled conditions and monitored for symptoms; infiltrated areas and systemic leaves were collected for RNA extraction using ‘Direct-zol RNA Kits’ (Zymo research, Irvine, CA, USA). Total RNA was DNase-treated, reverse transcribed and the replication of TsvlRNA1 was detected by minus-strand-specific RT-qPCR.

## Results

### *Trichoderma spirale* harbors an infectious circular viroid-like RNA

Pagnoni et al. (36) reported the identification of an orphan contig of circa 1.4 kb designated as ORFan2 in *T. spirale* (T45; see Supporting Information and Fig S1 for details on the identification of all isolates used in this study), which was isolated from an uncultivated soil in Sardinia. Recent bioinformatic analysis indicated that this contig corresponds to a putative circular RNA carrying a HHrbz in one polarity strand (Fig 2). The monomeric molecule of 712 nucleotides has been renamed Trichoderma spirale viroid-like RNA 1 (TsvlRNA1). In silico predictions using RNAfold indicated that the minimum free-energy secondary structure of TsvlRNA1 adopts compact rod-like and quasi rod-like conformation for the plus and minus polarity, respectively (Fig 2).

**Figure 2.**
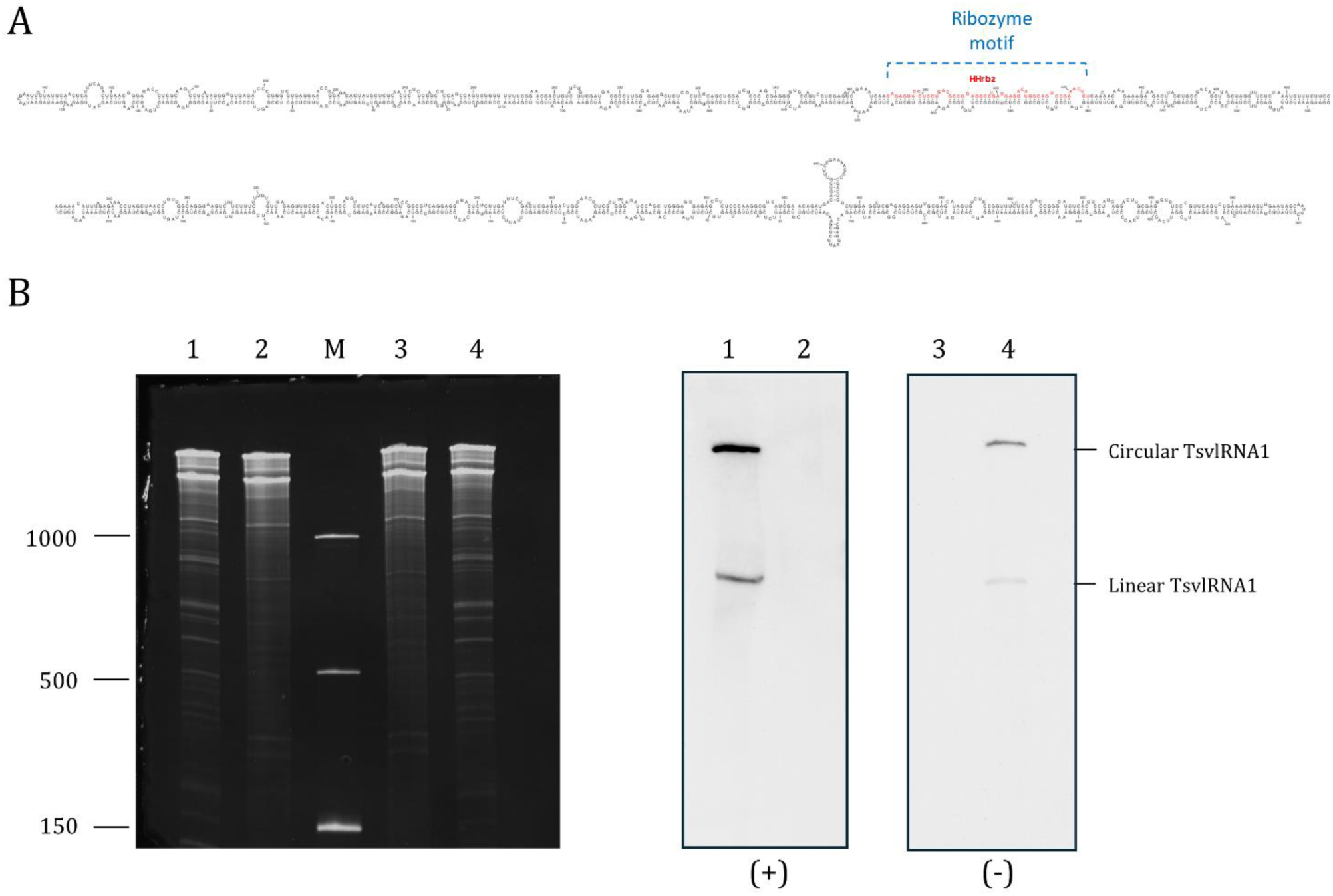
Main molecular features of TsvlRNA1. **A** Predicted secondary structure of lowest free energy calculated by RNAfold WebServer (http://rna.tbi.univie.ac.at/cgi-bin/RNAWebSuite/RNAfold.cgi) for the plus strand of Trichoderma spirale viroid-like RNA1. The region involved in the formation of the hammerhead ribozyme structure is delimited by a broken line. **B** Detection of circular and linear forms of both polarity strands of TsvlRNA1 by northern blot hybridization assays under denaturing conditions (5% PAGE in 8M urea 1X TBE). Lanes 1 and 4, RNA preparations from *Trichoderma spirale* isolate T45 tested positive to TsvlRNA1 by HTS and RT-qPCR; lanes 2 and 3 RNA preparations from the TsvlRNA1-negative *Trichoderma* isolate T43. M, RNA ssRNA Ladder (New England Biolabs) with RNA sizes (nt) indicated on the left. Identical aliquots of the same RNA preparations were loaded in parallel in the same PAGE (left, ethidium bromide staining of the gel). After nucleic acid transfer, the membrane was cut vertically (indicated with the broken white line) and each half was hybridized with equalized DIG-RNA probes specific to detect the plus (lanes 1 and 2) and the minus (lanes 3 and 4) polarity strands of TsvlRNA1 (right end panels). The positions of circular and linear forms of TsvlRNA1 are indicated on the right.

Reverse transcription with random primers followed by PCR using adjacent primers (Table S1) of opposite polarity produced an amplicon of the expected size, supporting the circular nature of TsvlRNA1. Northern blot analysis under denaturing conditions performed on *T. spirale* (T45) RNAs, revealed the presence of monomeric linear and circular forms of both polarity strands in vivo (Fig 2). This finding confirms the circular nature of TsvlRNA1 and supports its replication via a symmetric RCR mechanism, characteristic of members of the *Avsunviroidae* family. This replication variant is associated with two ribozymes, one on each polarity strand. However, TsvlRNA1 contains a known ribozyme solely in the plus strand —by convention, the most abundant in vivo. The self-cleaving activity of TsvlRNA1 RNAs of both polarity strands was assessed by in vitro transcription of a recombinant plasmid containing head-to-tail dimeric TsvlRNA1 cDNA insert, followed by analysis on denaturing 5% PAGE.

During transcription, monomeric and dimeric plus polarity TsvlRNA1 transcripts self-cleaved, generating fragments consistent with (+) HHrbz self-cleavage activity (Fig 3). Moreover, 5′ RACE experiments followed by cloning and sequencing confirmed that the 5’ termini of the 3′ cleavage fragment matched the predicted ribozyme cleavage site, confirming the specific processing site of the (+) HHrbz (Fig 3). In contrast, transcripts of the minus polarity strand lacking any identifiable ribozyme did not self-cleave during in vitro transcription, showing one band corresponding to the full-length RNA transcript (Fig 3). Attempts to induce post-transcriptional in vitro self-cleavage of purified dimeric (−) TsvlRNA1 transcripts by denaturation and slow renaturation under high Mg²⁺ and varied pH conditions were unsuccessful. These findings suggest either the existence of a yet undiscovered ribozyme with specific activation requirements or the involvement of a host fungal RNase in the replication of this vdlRNA.

**Figure 3.**
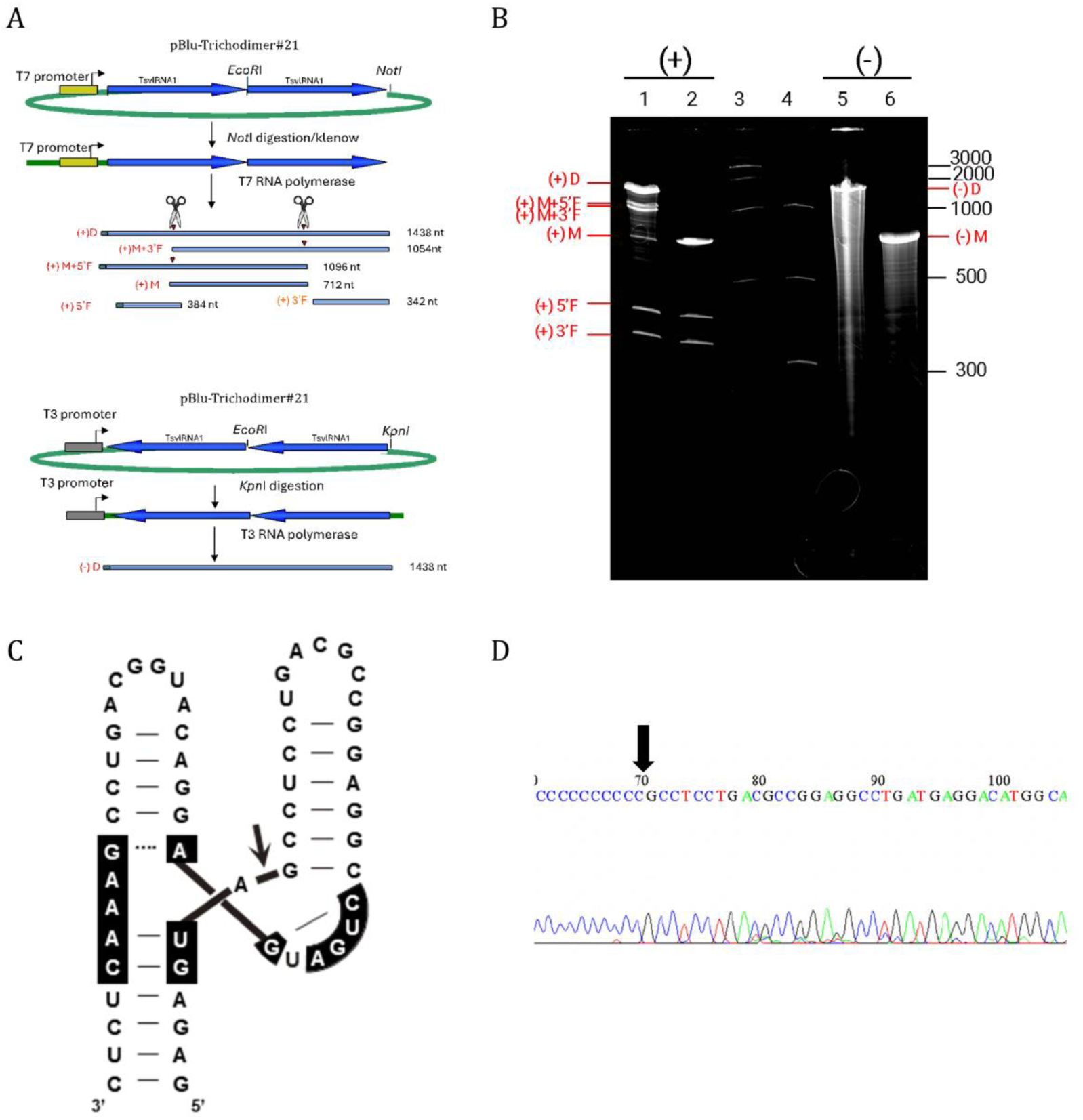
Self-cleavage activity in vitro of TsvlRNA1. **A** Schematic representation of the plasmid pBlu-Trichodimer#21, containing the dimeric head-to-tail cDNA sequence of TsvlRNA1, used as template for in vitro transcription and the expected RNA products with size reported on the right. Scissors mark the position of the self-cleavage sites. pBlu-Trichodimer#21 linearized with *Eco*RI and transcribed with T7 or T3 RNA polymerase produces monomeric RNAs (M) of (+) or (−) polarity strands, respectively. **B** Analysis by 5% PAGE under denaturing conditions of the in vitro transcription of plasmid pBlu-Trichodimer#21. Lanes 1, and 2, in vitro transcription of dimeric and monomeric (+) TsvlRNA1, respectively; lanes 5 and 6 in vitro transcription of dimeric and monomeric (−) TsvlRNA1, respectively; Lane 3, ssRNA ladder RNA (New England Biolabs); lane 4, low range ssRNA ladder (New England Biolabs). Sizes in nt are indicated on the right. In red are indicated the RNA fragments generated during transcription (see panel A). **C** Secondary structure of (+) hammerhead ribozyme of TsvlRNA1. The arrow indicates the predicted self-cleavage site and the conserved motifs in most HHrbz are denoted in black background. **D** Sanger sequencing electropherogram of the 5’ RACE product of the (+) 3’ fragment generated by (+) TsvlRNA1 HHrbz self-cleavage during in vitro transcription. The 5′ terminal nucleotide is indicated by an arrow.

### TsvlRNA1 replication is initiated from the in vivo transcription of dimer viroid-like cDNA

We initially engineered a monomer and head-to-tail dimer cDNA of TsvlRNA1 in the fungal expression vector pXH9 (Fig 1). Transformants from isolates *T. lixii* (T36), *T. spirale* (T45neg) and *T. atrobrunneum* (T99) showing mitotic stability for hygromycin B resistance were analyzed to confirm transgene integration through qPCR using TsvlRNA1-specific primers (Table S1) on total nucleic acid samples.

Northern blot analysis using probes for both plus and minus polarities was performed on DNase-treated RNA samples from the transformants. The analysis revealed the presence of monomeric linear and circular forms of both polarities in the ‘pXH9 dimer’ but not in the ‘pXH9 monomer’ transformants (Fig 4). These results provide evidence of replication of TsvlRNA1 and support that its replication is initiated only in the presence of the dimeric transgene generating the genomic monomeric linear forms by self-cleavage (Fig 1).

**Figure 4.**
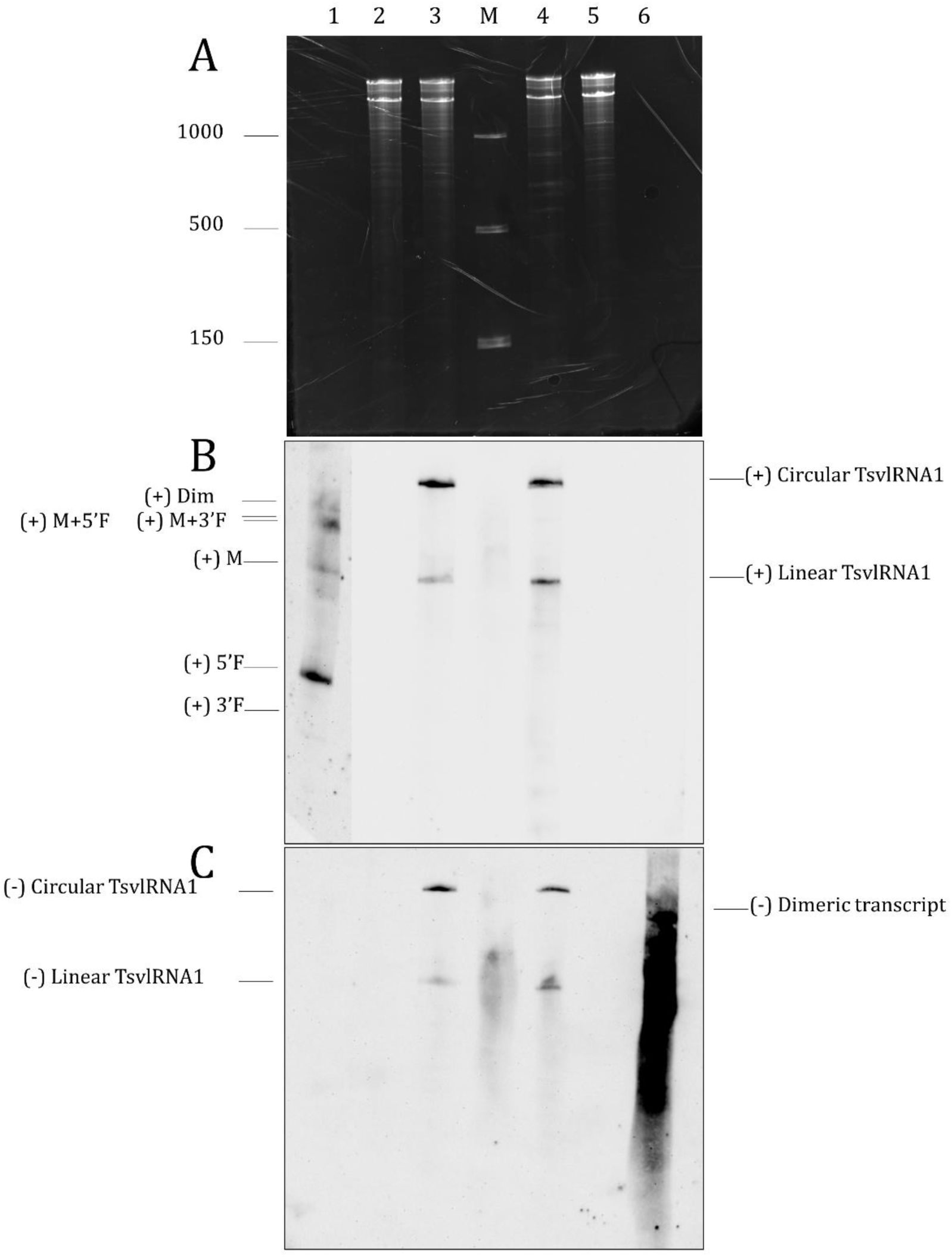
Detection of circular and linear forms of both polarity strands of TsvlRNA1 by northern blot hybridization in transgenic fungal isolates. Assays were performed under denaturing conditions (5% PAGE in 8M urea 1X TBE). Lane 1 in vitro transcription of dimeric (+) TsvlRNA1, lane 2 RNA preparations from the untransformed and TsvlRNA1-negative *Trichoderma atrobrunneum* isolate T99, lane 3 RNA preparations from *T. spirale* isolate T45 which tested positive to TsvlRNA1 by HTS and RT-qPCR, lane 4 RNA preparations from *T. atrobrunneum* isolate T99 transformed with ‘pHX9 dimer’, lane 5 RNA preparations from *T. atrobrunneum* isolate T99 transformed with ‘pHX9 monomer’, and lane 6 in vitro transcription of dimeric (−) TsvlRNA1. M, RNA ssRNA Ladder (New England Biolabs) with RNA sizes (nt) indicated on the left. Identical aliquots of the same RNA preparations were loaded in parallel in the same PAGE. **A** Ethidium bromide staining of the gel. After nucleic acid transfer, the membrane was hybridized with equalized DIG-RNA probes specific to detect the plus (panel **B**) and the minus (panel **C**) polarity strands of TsvlRNA1. The positions of circular and linear forms of TsvlRNA1 are indicated.

However, this does not apply to *T. lixii* (T36); although the transformation was successful, minus strand-specific RT-qPCR negative results revealed the absence of TsvlRNA1 replication (Table 2).

**Table 2.** Table summarizing the quantification cycle (Cq) values resulting from Real-Time qPCR analysis conducted on samples of DNase-untreated total nucleic acid, DNase-treated RNA, cDNA synthesized with random primers from DNase-treated RNA, and cDNA synthesized with minus sense-specific primer from DNase-treated RNA. RNA samples were extracted from protoplasts of *T. lixii* (T36), *T. spirale* (T45), and *T. atrobrunneum* (T99) transformed with the plasmids ‘pHX9 monomer’, ‘pHX9 dimer’, ‘pHX9 RBZ-A’, ‘pHX9 RBZ-C’, and ‘pHX9 RBZ-T’. For each sample, the mean of the two Cq values obtained from the technical replicates is reported. Samples are considered positive when Cq < 30 and are highlighted in yellow.

| Transformants | Total nucleic acid | DNase-treated RNA | cDNA | Minus Strand |
| --- | --- | --- | --- | --- |
| T99 DIM 3 | 25.5 | 36.4 | 24.2 | 26.8 |
| T99 DIM 25 | 25.6 | 38.1 | 24.9 | 26.4 |
| T99 DIM 52 | 25.8 | ND | 24.4 | 26.4 |
| T99 MONO 15 | 25.9 | ND | 36.5 | 35.9 |
| T99 MONO 24 | 26 | ND | 35 | 34.9 |
| T99 MONO 26 | 25.9 | ND | 34.2 | 34.1 |
| T45 DIM 5 | 24.6 | 34.1 | 22.9 | 22.6 |
| T45 DIM 7 | 24.6 | 33.5 | 21 | 22.1 |
| T45 DIM 14 | 20 | 30.4 | 21.7 | 21.6 |
| T36 DIM 39 | 23.6 | 35.6 | 31 | ND |
| T36 DIM 41 | 22.8 | 35.2 | 30.3 | 38.6 |
| T36 DIM 46 | 22 | 34.1 | 30 | 37.2 |
| T99 RBZ -ΔA 17 | 26.2 | 33.2 | 33.8 | 35 |
| T99 RBZ -ΔA 18 | 26.2 | 33.5 | 32.9 | 38.4 |
| T99 RBZ -ΔA 33 | 21.9 | 30.6 | 31.1 | 34.3 |
| T99 RBZ T 2 | 26.5 | 33.4 | 33.3 | ND |
| T99 RBZ T 27 | 25.6 | 32.1 | 32.1 | 36.4 |
| T99 RBZ T 31 | 28.3 | 32.3 | 31.2 | ND |
| T99 RBZ C 21 | 27.1 | 32.6 | 34 | 35.8 |
| T99 RBZ C 26 | 26.1 | 32.6 | 33 | 38.7 |
| T99 RBZ C 29 | 24.8 | 31.9 | 31.9 | 35.3 |

### Horizontal transmission of TsvlRNA1

A critical test for establishing vdlRNAs infectivity is their capacity to disseminate throughout the host population. In general, horizontal transmission of mycoviruses occurs through anastomosis (hyphal fusion). We therefore conducted experiments using dimer-transformed viroid-like isolates and viroid-like-free recipient isolates (all combinations are shown in Table 1).

The presence of TsvlRNA1 in recipient strains was confirmed through RT-qPCR and minus strand-specific RT-qPCR using as template cDNA synthesized from RNA extracted from colonies unable to grow on hygromycin B-containing medium (Table S3). When *T. atrobrunneum* (T99DIM3) transformed with the plasmid ‘pXH9 dimer’ was the donor strain, TsvlRNA1 was transmitted to *T. afroharzianum* (T22; TO71B), *T. gamsii* (T71), *T. atrobrunneum* (T84; T99); when *T. spirale* transformed with the same plasmid (T45DIM5) was the donor, transmission occurred to *T. spirale* (T45neg), *T. gamsii* (T71), *T. atrobrunneum* (T84), and *T. velutinum* (T100). Transmission failures were observed between certain donor-recipient pairs (Table 1). Northern blot analysis confirmed monomeric linear and circular forms of both strands in recipient strains, demonstrating that TsvlRNA1 is a replicative and infectious RNA (Fig 5). These results show that TsvlRNA1 can spread horizontally in *Trichoderma* species, resulting in isogenic, transgene-free infected colonies, and provide the first evidence of transmission to species beyond its original host, *T. spirale*. None of the fungal species’ pairings tested for TsvlRNA1 transmission showed evidence of barrage (incompatible reactions), supporting the possibility that interspecific TsvlRNA1 transmission relies on anastomosis of compatible hyphal fusion among different species (see Fig S2 and Supporting Information for details).

**Figure 5.**
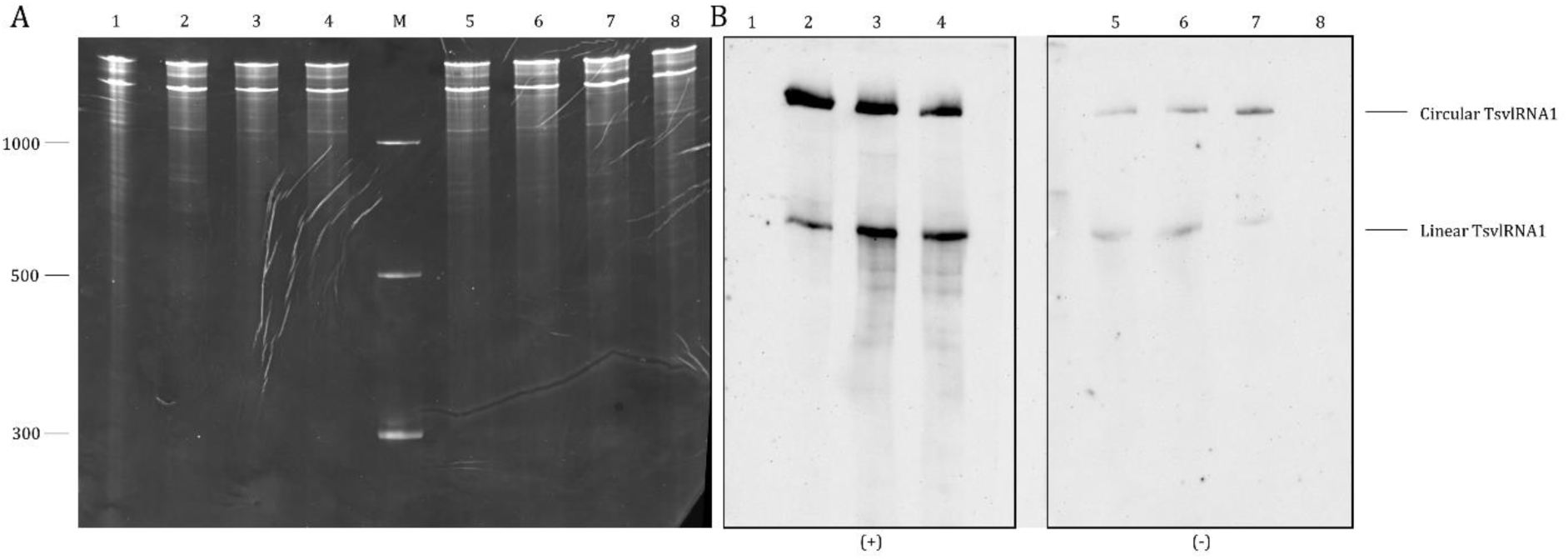
Detection of circular and linear forms of both polarity strands of TsvlRNA1 in horizontally transfected isolates by northern blot. Hybridization assays were carried out under denaturing conditions (5% PAGE in 8M urea 1X TBE). Lanes 1 and 8 RNA preparations from the TsvlRNA1-negative *Trichoderma spirale* isolate T45neg, lanes 2 and 7 RNA preparations from *T. spirale* isolate T45 which tested positive to TsvlRNA1 by HTS and RT-qPCR, lanes 3 and 6 RNA preparations from *T. spirale* isolate T45DIM5 transformed with ‘pHX9 dimer’, lanes 4 and 5 RNA preparations from *T. spirale* isolate T45ANAS-A obtained through horizontal transmission. M, RNA ssRNA Ladder (New England Biolabs) with RNA sizes (nt) indicated on the left. Identical aliquots of the same RNA preparations were loaded in parallel in the same PAGE. **A** Ethidium bromide staining of the gel. After nucleic acid transfer, the membrane was cut vertically (indicated with the broken white line) and each half was hybridized with equalized DIG-RNA probes specific to detect the plus (lanes 1, 2, 3, and 4) and the minus (lanes 5, 6, 7, and 8) polarity strands of TsvlRNA1 (Panel **B**). The positions of circular and linear forms of TsvlRNA1 are indicated on the right.

Based on prior virome characterization (36), some of the isolates we are using harbor mycoviruses (Table S4). Nevertheless, successful horizontal transmission of TsvlRNA1 from the virus-free donor strain of *T. spirale* (T45DIM5) to the virus-free recipient strains *T. spirale* (T45neg), *T. gamsii* (T71), and *T. velutinum* (T100) indicates that, like viroids, TsvlRNA1 replication does not rely on a helper virus. Although T84 and T99 are both classified as *T. atrobrunneum*, TsvlRNA1 transmission from T45DIM5 was detected only in T84, suggesting that strain-level differences, may affect TsvlRNA1 transmissibility (Table S3). When T99DIM3 was used as the donor, the present mycoviruses were co-transmitted together with the vdlRNA to different *Trichoderma* species (Table S4).

### Vertical transmission and acquisition of environmental RNA do not represent modes of dissemination for TsvlRNA1 among *Trichoderma* isolates

Vertical transmission of TsvlRNA1 was assessed in conidial progeny of infected isolates. Thirty-two single-conidium-derived colonies from TsvlRNA1-positive *T. atrobrunneum* isolates (T84ANAS-1 and T99ANAS-1), obtained via horizontal transfer, were analyzed. RT-qPCR revealed that none of the progeny carried TsvlRNA1. All colonies derived from T99 hyphal fragments exposed to total RNA from TsvlRNA1-positive T45ANAS-A or to viroid transcripts synthetized in vitro tested negative in northern blot analysis. These results indicate that TsvlRNA1 is neither vertically transmitted via conidia nor acquired from environmental RNA.

### Infectious clone mutants of TsvlRNA1

To evaluate the functional relevance of the (+) HHrbz of TsvlRNA1 in its autonomous replication, targeted mutagenesis was employed to affect ribozyme self-cleavage.

Protoplasts from *T. atrobrunneum* (T99) were transformed with plasmids ‘pHX9 RBZ -ΔA’, ‘pHX9 RBZ C’, and ‘pHX9 RBZ T’, which carry a nucleotide deletion or a single point mutation in the HHrbz catalytic core (Fig 1). Although transgene integration was confirmed by qPCR, no TsvlRNA1 replication was detected by minus strand-specific RT-qPCR (Table 2). These results demonstrate that a catalytically active ribozyme is essential for TsvlRNA1 replication.

### RNAi response to TsvlRNA1

sRNAs profiles from TsvlRNA1 were compared with those of co-infecting mycoviruses across multiple *Trichoderma* isolates, including two isolates harboring cytoplasmic mycoviruses (T22ANAS-1 and T99ANAS-1) and two virus-free isolates (T45ANAS-A and T71ANAS-A). Viroid-like-derived small RNAs (vd-sRNAs) were mainly 20–23 nt long, with 21-nt species predominating, closely resembling virus-derived sRNA profiles (Fig S3). Plus-strand sRNAs were slightly more abundant than minus-strand sRNAs, consistently with the preferential accumulation of plus polarity RNA species in the host (Fig S4). Mapping of TsvlRNA1-derived sRNA reads identified several hotspots without clear functional association: between nucleotides 47 and 67 on the plus strand, and between nucleotides 620 and 647 on the minus strand (Fig S3). The predominant 21-nt class showed enrichment for adenine (A) or cytosine (C) at the 5′ terminus, with no strand-specific differences.

### Effect of TsvlRNA1 infection on antagonistic capacity of *Trichoderma* species

Since *Trichoderma* genus is commonly used as a biocontrol agent in agriculture, different pairs of TsvlRNA1-infected and TsvlRNA1-free isogenic *Trichoderma* strains obtained from co-culture (T45ANAS-A and -C, T71ANAS-2, TO71BANAS-1 and -2, T84ANAS-1 and -A, and T99ANAS-1) were evaluated for their capability to reduce the damage caused by *R. solani* on small broad bean seedlings. Although several strains harbored multiple mycoviruses (Table S4), complicating attribution of biological effects exclusively to the vdlRNA element, results consistently suggested that TsvlRNA1 infection modulates antagonistic activity (Fig S5). Among the cases in which the specific contribution of the vdlRNA could be more clearly discerned, TsvlRNA1 significantly enhanced antagonism in *T. atrobrunneum* (T99ANAS-1) against pre- and post-emergence damping-off (Fig 6A), whereas in *T. spirale* (T45ANAS-A and -C) the effect on antagonistic activity was negative (Fig 6B).

**Figure 6.**
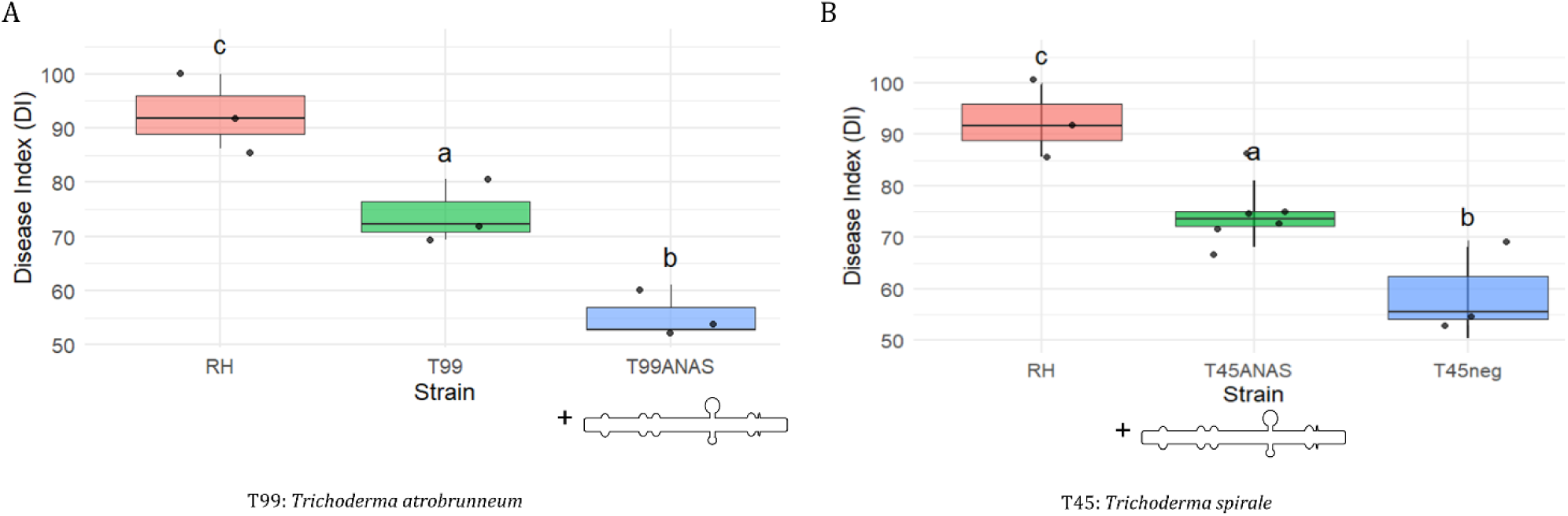
Box plots illustrate the evaluation of *Trichoderma* antagonistic activity against *Rhizoctonia solani* infection in small broad beans. Box plots illustrate the assessment of *Trichoderma* antagonistic activity against *Rhizoctonia solani* infection in small broad beans. In the box plots corresponding to isolates obtained by co-culture (T45ANAS-A and -C, T99ANAS-1), the data points reflect the aggregation of multiple biological and experimental replicates. For *R. solani* control (RH) and the original recipient isolates (T45neg and T99), the data derives from three biological replicates. Statistical analyzes were performed using one-way ANOVA followed by Honestly Significant Difference post-hoc test and the Kruskal–Wallis test followed by Dunn’s post-hoc test. Strains infected by the TsvlRNA1 are represented by a graphical symbol under the name of the isolate in the X axis. **A** A positive effect of TsvlRNA1 presence on the ability of *T. atrobrunneum* (T99ANAS-1) to limit the *R. solani* caused symptoms compared to the same strain without TsvlRNA1 (T99). **B** Reduction of the antagonistic capacity for strains of *T. spirale* with TsvlRNA1 presence (T45ANAS-A and -C) to limit the *R. solani* symptoms. DI: Disease index. Strain sharing the same letters are not statistically different.

### No evidence of TsvlRNA1 replication in plant cell

The same monomeric and dimeric TsvlRNA1-cDNA molecules used for *Trichoderma* experiments were cloned between the 35S promoter and terminator of the binary vector pJL22 for expression in *N. benthamiana* via *A. tumefaciens*. Seven days post agroinfiltration, no significant differences in plus-strand accumulation were detected between the constructs, and minus strand accumulation was undetectable for both constructs in infiltrated areas and systemic leaves (Table S5). Local complementation with TMV movement protein did not affect RNA accumulation.

## Discussion

This study reports the first in silico identification of a vdlRNA, called TsvlRNA1, in the non-phytopathogenic fungus *T. spirale*. Diener’s proposal that viroids might represent relics of the primordial RNA world was regarded with skepticism, as confirmed hosts were restricted to plants and later to animals (49,50). Recent metatranscriptomic studies, including this work, have reshaped this perspective by identifying fungal vdlRNAs, comprising ambiviruses and mitoviruses (5,51,52), and Obelisks in bacteria (22). Nevertheless, assessing common ancestry of these agents remains challenging due to their low level of sequence conservation (52).

The circular nature of TsvlRNA1 is supported by PCR amplification of a full-length product with adjacent primers of opposite polarity, northern blot detection of circular monomers of both polarities in vivo, and bioinformatic identification of direct repeated sequences. Although the detection of circular and linear RNA monomers of both polarities is indicative of a symmetric RCR mechanism involving ribozymes in both polarity strands (2), only a single HHrbz in the plus strand of TsvlRNA1 was predicted and experimentally validated in vitro, while no ribozyme or cleavage activity was detected in the minus strand. This represents the first example of a vdlRNA undergoing symmetric RCR while producing circular RNAs of both polarities despite having a ribozyme in only one strand. Although the vdlRNAs described by Dong et al. (51) from *Botryosphaeria dothidea* accumulate circular and linear forms of both strands, consistent with our observations, they lack recognizable ribozymes, and only one of them exhibits in vitro self-cleavage of one strand. This finding deviates from the canonical symmetric RCR model and suggests a replication strategy partially resembling that of certain viroid-like satellite RNAs (e.g. *Solanum nodiflorum* mottle virus and subterranean clover mottle virus) (15), or likely involving an as-yet unidentified ribozyme or a fungal host-derived RNase. This mechanistic discrepancy remains unresolved and warrants further investigation. To establish a reverse genetic system, infectious cDNA clones of TsvlRNA1 were generated, representing, to our knowledge, the first fungal vdlRNA clone with an experimentally validated ribozyme. Head-to-tail dimeric construct was essential for replication, whereas monomeric sequence was non-infectious, consistent with plant viroid studies (53). Replication was confirmed via RT-qPCR and northern blot analysis, and horizontal transmission to cDNA-free strains demonstrated autonomous replication independent of the transgene and/or helper virus. Mutant infectious clones affecting TsvlRNA1 HHrbz catalytic activity demonstrated that a functional ribozyme is essential for replication.

TsvlRNA1 can horizontally transmit and replicate autonomously across different *Trichoderma* species, independent of co-infecting mycoviruses, as demonstrated by transferring from a virus-free donor to three different virus-free recipient species. These results indicate that TsvlRNA1 is not a viroid-like satellite RNA, but rather an infectious agent endowed of autonomous replication within the infected cells. Considering that mycoviruses are typically transmitted via hyphal fusion between isolates of the same VCG, and that intraspecies transmission is restricted by VIC (31), the observed horizontal transmission of TsvlRNA1 and other viruses among different *Trichoderma* species belonging to distinct sections —taxonomic units comprising evolutionarily related species separated by substantial genetic and temporal distances— is noteworthy (54). Moreover, to the best of our knowledge, no studies have specifically addressed interspecific interactions among the species examined here. It is plausible that previous studies have overestimated the ability of VIC to inhibit virus transmission, as on-site investigations indicate that mycoviruses can spread efficiently among vegetatively incompatible strains, suggesting that environmental factors may facilitate transmission (55). Despite evidence that extracellular RNAs can persist in a stable and biologically active form (56,57), our results indicate that environmental acquisition of TsvlRNA1 does not represent a mode of transmission. Therefore, additional transmission mechanisms, such as the secretion and uptake of extracellular vesicles, cannot be excluded. As vertical transmission was not detected, horizontal transmission appears to represent the predominant mode of dissemination for TsvlRNA1, plausibly enabling its long-term persistence within natural fungal populations.

Among the *Trichoderma* isolates tested, *T. lixii* (T36) was the only strain in which TsvlRNA1 replication was undetectable, despite successful integration of the dimeric transgene. Furthermore, vdlRNA transmission to T36 did not occur from either *T. spirale* (T45DIM5) or *T. atrobrunneum* (T99DIM3). This resistance may reflect differences in host factors required for TsvlRNA1 replication.

RNA silencing represents a eukaryotic defense against invasive nucleic acids, triggered by double-stranded RNAs (dsRNAs) or structured single-stranded RNAs (ssRNAs) and mediated by Dicer-like ribonucleases (58). Viroids are strong inducers of this response due to their circular, highly base-paired ssRNA genome and replication via dsRNA intermediates (59). We provide the first analysis of sRNA populations in fungi infected by a vdlRNA. All TsvlRNA1-positive isolates accumulated 20–23 nt sRNAs with a dominant 21-nt peak, regardless of co-infection with other mycoviruses. This profile resembles sRNA patterns observed during infection by cytoplasmically replicating mycoviruses (60) and chloroplast-replicating viroids of the *Avsunviroidae* family (61), which are characterized by the accumulation of 19–22 nt sRNAs, peaking at 21 nt, and of 21–22 nt sRNAs, respectively. This reflects a conserved RNA silencing response across diverse eukaryotic lineages.

*Trichoderma* spp. are widely used biocontrol agents (26) and the investigation of their antagonistic capacity against *R. solani*, a destructive necrotrophic plant pathogen, is of major interest. Antagonistic trials using isogenic *Trichoderma* strains revealed that TsvlRNA1 can modulate biocontrol efficacy either positively or negatively in a strain/species-dependent manner; moreover, in most tested strains, co-infecting mycoviruses hindered the specific effect of TsvlRNA1. Enhanced antagonism in *T. atrobrunneum* (T99ANAS-1) suggests potential applications in fungal bioformulations, whereas the negative effect in *T. spirale* (T45ANAS-A and -C) highlights risks associated with field deployment, particularly due to possible horizontal transmission to native species and unpredictable outcomes.

The proven phenotypic effect of the vdlRNA alone, or in interactions with mycoviruses, suggest that searching for such elements through well-established bioinformatic pipelines should be a gold standard in studies that associate genotype to phenotype. These elements are functional components of fungal microbiomes and can be lost or gained experimentally when evaluating other fungal phenotypic features (site directed mutagenesis, mycovirus transfection/curing, etc.). Our study implies that ribozyme detection in a single orientation should be included in vdlRNA detection bioinformatic pipelines, increasing the number of vdlRNAs previously overlooked.

Our findings on horizontal transmission among different *Trichoderma* species are somewhat surprising since they encompass both vdlRNA and mycoviruses. Only occasional indirect evidence has been reported in the literature regarding the isolation of the same viral species from different fungal hosts, in one case belonging to distinct classes (62–65). Moreover, interspecific transmission in in vitro co-culture experiments has also been sporadically documented (64,66,67). To further support our findings from controlled-environment experiments, we noticed that in the *Trichoderma*-associated virome dataset reported by Pagnoni et al. (36), several mycoviruses were detected in different *Trichoderma* species originating from the same geographical region (Table S6). Given the importance of horizontal transmission of cytoplasmic elements for resistance to antifungals (a sanitary emergency world-wide) (68,69), we propose to establish a complex interspecific *Trichoderma* hyphal network monitoring mycovirus and vdlRNA transmission as an experimental system to study the mechanism of interspecific hyphal fusion. It is worth speculating that the mycoparasitic lifestyle of *Trichoderma* spp. may confer a specific propensity for hyphal fusion, potentially bypassing the pre- and post-fusion checkpoints characterized in other genera within the Pezizomycotina subphylum (70,71).

## ACKNOWLEDGEMENTS

The authors performed experiments at the Imaging Platform of the Department of Translational and Molecular Medicine of the University of Brescia.

## Abbreviations

(ccc): covalently closed circular
(HHrbz): hammerhead ribozymes
(RCR): rolling-circle replication
(vdlRNA): viroid-like RNA
(VIC): vegetative incompatibility
(VCG): vegetatively compatible groups
(TsvlRNA1): Trichoderma spirale viroid-like RNA 1
(RT-qPCR): Real-Time qPCR
(*gpd-1*): glyceraldehyde-3-phosphate dehydrogenase
(sRNA): small RNA
(vd-sRNA): viroid-derived small RNA
(dsRNA): double-stranded RNA
(ssRNA): single-stranded RNA
(CHV1): Cryphonectria hypovirus 1
(SRA): Sequence Read Archive
(HTS): High-Throughput Sequencing
(PAGE): polyacrilammide gel electrophoresis
(PDA): potato dextrose agar
(ITS): internal transcribed spacer
(RPB2): RNA polymerase II subunit
(TEF1-α): translation elongation factor 1-α gene
(MIC): minimum inhibitory concentration
(GFP): green fluorescent protein
(TMV): tobacco mosaic virus

## Ethics and Integrity policy

### Data availability statement

The resulting raw sequencing reads have been deposited in the NCBI Sequence Read Archive (SRA) under BioProject PRJNA1303385 (Accession Number SRR34918256), PRJNA1253966 (Accession Number SRR33245418 and SRR33245416), and PRJNA1301113 (Accession Number SRR34855465, SRR34855464, SRR34855463, and SRR34855462).

## Supporting Information

Supporting Information containing further Text, Figures and Tables is available as a link

